# Reduced childhood social attention in an autism model marmoset predicts impaired social skills and inflexible behavior in adulthood

**DOI:** 10.1101/2022.03.03.482182

**Authors:** Akiko Nakagami, Miyuki Yasue, Keiko Nakagaki, Madoka Nakamura, Nobuyuki Kawai, Noritaka Ichinohe

**Affiliations:** Japan Women’s University; National Center of Neurology and Psychiatry; Nagoya University

**Keywords:** early intervention, gaze direction, valproic acid, restricted behavior, third-party reciprocity, inequity aversion, reversal learning, longitudinal study

## Abstract

Autism spectrum disorder (ASD) is a neurodevelopmental disorder characterized by social and communication impairments and restricted and repetitive behavior. Although currently no established cure exists for ASD, early intervention to the deficits of attention to other individuals is expected to reduce the progression of ASD symptoms in later life. In order to confirm this hypothesis and improve early therapeutic interventions, it is desirable to develop an animal model of ASD in which social attention is impaired in childhood and ASD-like social behavior is observed in adulthood. However, rodent models of ASD have difficulty in recapitulating the deficit of gaze-based social attention. In this study, we examined the direction of gaze towards other conspecifics during childhood and puberty in a three-chamber test setting using an ASD model of marmoset produced by maternal exposure to valproic acid (VPA). We also conducted a reversal learning test in an adult VPA-exposed marmoset as an indicator of perseveration, a core symptom of ASD that has not previously been investigated in this model. The results showed that time spent gazing at other conspecifics was reduced in VPA-exposed marmosets in childhood, and adults displayed rigidity of response. In a longitudinal study using the same animals, deficits in social attention in childhood correlated well with ASD-like social disturbance (inequity aversion and third-party reciprocity) and inflexible behavior in adulthood. Since VPA-exposed marmosets exhibit these diverse ASD-like behaviors that are coherent from childhood to adulthood, VPA-exposed marmosets will provide a valuable opportunity to elucidate mechanisms for early intervention and contribute to the development of early therapies.

## Introduction

Autism spectrum disorder (ASD) is a neurodevelopmental disorder characterized by social and communication impairments and restricted or repetitive behaviors (1). The prevalence of ASD in children has been increasing over the years, and one in 44 children is currently diagnosed with ASD (2). However, at this time, there is no established cure for ASD. Research has shown that early interventions for ASD are more likely to have major long-term positive effects on symptoms and later skills (3,4), and initiating interventions is recommended as soon as a diagnosis of ASD is seriously considered or determined (4–7). A hypothesis for the mechanism of the early treatment effect is that the prodrome of ASD accelerates the development of later symptoms (8–11). In particular, early deficits in attention to the eyes and faces of others are thought to interfere with normal social input that promotes the proper development of social brain circuits (3,11,12). Furthermore, it is suspected that, after early brain plasticity is reduced, the aberrantly formed circuits will be difficult to reverse. In order to confirm this set of hypotheses and to establish effective therapeutic interventions, development of animal models of ASD with impaired social attention in childhood and higher order ASD-like behavior in adulthood is desirable.

Rodent models of ASD have provided much insight into ASD to date. However, rodent models have difficulty in recapitulating the deficit of gaze-based social attention and impairments in higher-order social skills. In addition, neurodevelopmental events between birth and weaning in rodents occur between mid-gestation and birth in primates (13), making it harder to match the age of human childhood in rodents. The common marmoset (*Callithrix jacchus*), the New World monkey, is a promising new model for ASD because of its rich social skills, including eye contact, vocal communication, and evaluation of third-party reciprocity (14–17). The early social environment of marmosets influences later social skill acquisition. For example, the interaction of marmoset offspring with their parents influences the development of the acoustic properties of their calls, advanced communicative behaviors such as turn-taking, and social play (18–21). Furthermore, marmosets are easy to handle, mature rapidly, and have high reproductive capacity. These characteristics suggest that the marmoset can be used to study the treatment of ASD through early modulation of social attention. We established an ASD marmoset model by administering valproic acid (VPA) *in utero* (22–24). VPA modifies gene expression in the developing fetal brain by inhibiting histone deacetylase (HDAC). Marmosets exposed to VPA showed atypical calls in childhood (24). As adults, VPA-exposed offspring could not qualify for advanced social testing (22,23), as they were unable to recognize whether or not another individual reciprocated (22). Inequity aversion, which is hypothesized to have evolved in conjunction with cooperation (25), was not found in VPA-exposed marmosets (23), similar to individuals with ASD (26,27). Moreover, the transcriptome of the marmoset cortex model has been shown to reproduce the features of human ASD better than rodent models (24). Morphological analysis of VPA-exposed marmosets has revealed insufficient commissural fibers and altered immune cells in the brain, similar to human ASD (28,29).

In the present study, we performed a three-chamber social test on VPA-exposed marmosets from childhood through puberty to examine deficits in social attention. In addition to the chamber preference of the model animals, the direction of gaze to other conspecifics was assessed. We also conducted a reversal learning test in an adult model marmoset as an indicator of preservation, a core symptom of ASD that has not been previously investigated in this animal model. The results showed that VPA-exposed marmosets exhibited reduced social attention in childhood and that adult animals displayed rigidity of response. In a longitudinal study using the same animals, social attention deficit in childhood correlated well with ASD-like social disturbance (inequity aversion and third-party reciprocity) (22,23) and inflexible behavior in adulthood. Since VPA-exposed marmosets exhibit these diverse ASD-like behaviors that are coherent from childhood to adulthood, VPA-exposed marmosets will provide a valuable opportunity to elucidate mechanisms for early intervention and contribute to the development of early therapies.

## Materials and Methods

All experimental and animal care procedures were performed in accordance with NIH guidelines and the Guide for Care and Use of Laboratory Primates published by the National Institute of Neuroscience, National Center of Neurology and Psychiatry, and were approved by the Animal Research Committee at the National Institute of Neurosciences in Tokyo, Japan. Marmosets were bred in the National Center of Neurology and Psychiatry, kept under a 12 h/12 h light/dark cycle, and provided food (CMS-1, CLEA Japan) and water ad libitum. Temperature and humidity were maintained at 27–30 °C and 40%–50%, respectively. To produce marmosets prenatally exposed to VPA, serum progesterone levels in the female marmosets were monitored once a week to determine the timing of pregnancy. We took a minimum amount of blood (<0.1 mL) for the assay and provided the marmoset with extra nutrition after blood collection. In addition to the blood progesterone level, pregnancy was further confirmed by palpitations and ultrasound monitoring (Ultrasound Scanning; Xario, Toshiba Medical Systems Corp., Tochigi, Japan). We administered 200 mg/kg of sodium valproate (VPA, Sigma–Aldrich, St. Louis, MO, USA) seven times from day 60 to 66 after conception. The dose was calculated from the maximum dose used in human patients (1200 mg/day) by considering the difference in body surface area between marmosets and humans. We also considered the time of VPA administration to be comparable to embryonic day 12 (E12) in rodent models, the period in which VPA induction produced the most prominent abnormality in social behavior without any physical deformity (30). To avoid vomiting, marmosets were first given bread, and VPA was delivered to the stomach 30 min later through a 4-Fr feeding tube (Atom Medical Corp, Tokyo, Japan) placed through the mouth, down the esophagus, and into the stomach. Using this procedure, we did not observe obvious malformations or deformities in VPA-exposed marmosets. In this study, eleven UE (unexposed) marmosets (three male and eight females) and seven VPA (prenatal exposure to VPA) marmosets (one male and six females) ranging from 15 weeks to 2.2 years of age were used.

### Three-chamber test

Six VPA-exposed marmosets (one male and five females) and five UE marmosets (one male and four females) were subjected to a three-chamber test during childhood (15-19 weeks of age) and puberty (29–41 weeks). During each period, the experiments were conducted once a week, five times per individual. The apparatus for the three-chamber test comprised a rectangular three-chamber box (88 cm W × 42 cm D × 22 cm H). The center chamber (25 cm × 40 cm) and the side chambers (30 cm × 40 cm) were separated by opaque 6-mm thick Plexiglas walls with a central opening (12 cm × 12 cm) that allowed free access to each chamber (Figure 1A). Opaque sliding doors were placed at these openings. The center chamber had an entrance gate (16 × 16 cm) with a transparent door. Subjects entered their carrying cages voluntarily, and they were taken into the experimental room. Then, they spontaneously entered the central chamber of the experimental box through the entrance gate. At this time, the opaque door between each chamber was closed. Thus, the experimenters had no contact with subjects. While the subject stayed in the center chamber, two identical transparent cylinders (15 cm in diameter and 20 cm in height), large enough to contain a single adult marmoset, were placed vertically inside the apparatus, one in each side chamber. The cylinders had removable lids with ventilation holes, and gel absorption sheets were placed below and above the cylinders to prevent the lid from opening. Either cylinder was invisible from the other side chamber, but was visible from the center chamber. One empty cylinder was immediately replaced with a cylinder containing an unfamiliar marmoset which had been prepared in a separate room. The subjects could not see the cylinder exchange. Both doors to the side chambers were then opened for the subject to explore the entire test apparatus for 10 min. After a 5 min acclimation in the center chamber, the door to both side chambers was opened.

**Figure 1.**
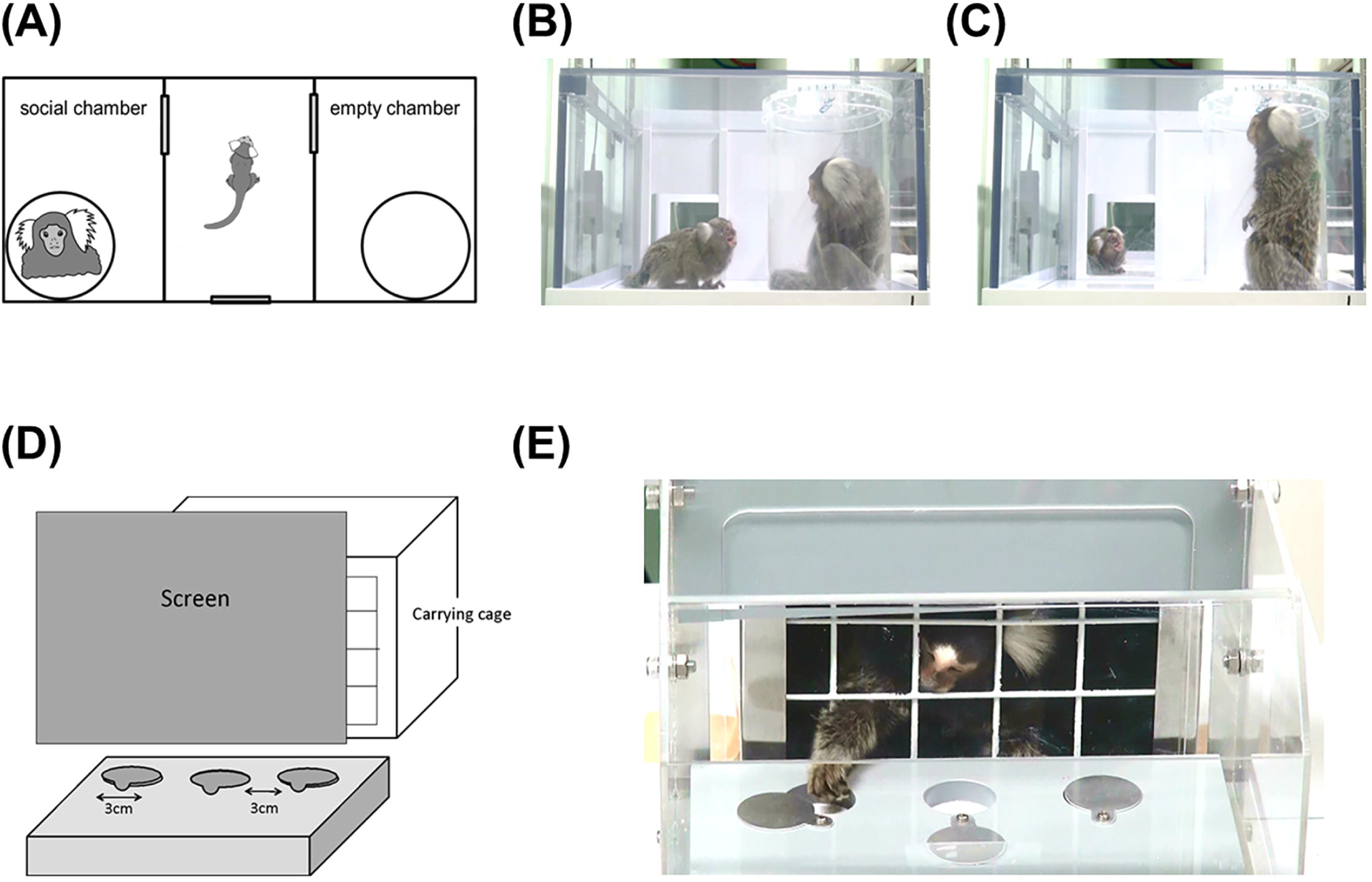
(**A**) Apparatus layout for the three-chamber test (88cm W × 42cm D × 22cm H). Three interconnected chambers are separated by two manually operated sliding doors. An unfamiliar adult marmoset was placed in one of the side chambers, stored in a clear plastic tube with air holes on the lid. Another empty tube was placed in the other side chamber. At the test phase, a subject marmoset went into the center chamber voluntarily with the sliding doors closed. Both doors to the side chambers were then opened for the animal to explore the entire test apparatus. (**B-C**) Representative images of social gazing at each chamber. (**B**) The duration of social gazing in the social chamber was measured. (**C**) The duration of social gazing in the center chamber was measured. (**D**) Apparatus layout for the reversal learning test. The experiment was conducted in a Wisconsin General Test Apparatus (WGTA) for the marmosets. The stimulus tray of the stimulus presentation chamber contained three wells, 3 cm in diameter, in a row. (**E**) The subject marmosets acclimated to open one of the slidable covers on the two wells to pick up a hidden food reward.

Five male and five female adult marmosets served as unfamiliar conspecific animals and met with target animals of the same sex only once. These unfamiliar adults were not kin of the subject, and had no previous interaction with the target animals, which may have created slight possibilities to provide visual, auditory, and olfactory cues.

The duration of time the subject spent in each chamber (social chamber, center chamber, and empty chamber) and the time head-orienting to the unfamiliar conspecific were counted (Figure 1B-C). The time to orient the head to an unfamiliar conspecific is an important measure to evaluate their social interest, because social attention in primate species heavily relies on visual cues; the subject does not necessarily need to come close to the unfamiliar marmoset. In this study, we used the orientation of marmosets’ heads to infer their gaze towards other individuals. This is justified because marmosets have more restricted eye motility than Old World monkeys and humans, as reflected by their oculomotor range (marmosets, ∼10°; macaques, 40-50°; humans, ∼55°) (31–33). In addition, animals with smaller heads, such as marmosets, are assumed to use head rotation rather than eye rotation to determine gaze, because the exertion from head rotation is smaller than that for animals with larger heads (31). Behaviors were recorded using three video cameras. The top video camera (Sony, handycam), located 73 cm above the apparatus, recorded the entire apparatus. Video cameras on each side captured activity near the cylinder and the open space of the center chamber at a distance of 40 cm from the side chamber. All recordings were synchronized and analyzed using the Observer XT 11 software (Noldus Information Technology, Wageningen, the Netherlands).

The duration of each epoch between the onset and offset of the behaviors was calculated by two observers. We analyzed reliability using two methods: the Duration/Sequence method and the Frequency/Sequence method. The Duration/Sequence method calculates the extent to which the scored duration of a given event is comparable between observations. The statistics showed that the percentage of agreement between the observers was 92.17% and Cohen’s Kappa was 0.90. The Frequency/Sequence method calculates the extent to which the scored frequency of given events is comparable between observations; we programmed a tolerance window of 1 s. The statistics showed that the percentage of agreement between observers was 84.58% and Cohen’s kappa was 0.80.

### Position reversal learning

Twelve juvenile (1.5 to 2.2 years old) marmosets (2 male and 4 female UE marmosets plus one male and 5 female VPA-exposed marmosets) were used in the reversal learning task. The experiment was conducted using a modified Wisconsin General Test Apparatus (WGTA) for marmosets. The WGTA was composed of a stimulus presentation chamber (20.5 cm x 25.0 cm x 20.0 cm) attached to a carrying box (Figure 1D-E). The stimulus tray of the stimulus presentation chamber (23.0 cm x 19.5 cm) contained three wells (3 cm in diameter) with slidable covers in a row.

Prior to the test, each marmoset underwent pre-training. First, they were trained to enter the carrying cage (20.0 cm x 25.0 cm x 18.5 cm) voluntarily. The monkeys learned to open the slidable covers on the two wells to pick up a hidden food reward (a piece of sponge cake; 0.5 cm x 0.5 cm x 0.5 cm). The test was initiated when the marmoset could reliably obtain a food reward from the left or right presentation well.

In this study, instead of conventional reversal learning for marmosets in which the food position was reversed within the same day (34–36), we employed reversal learning in which the reward side was inverted in the session that took place after the day the criterion was reached. The purpose of this procedure was to assess long-term cognitive flexibility deficits and higher-order restricted and repetitive behaviors such as obsession with habit and sameness. The test consisted of three phases: the first phase of discrimination learning (Phase 1), first reversal learning phase (Phase 2), and second reversal learning phase (Phase 3). The task involved positional learning, in which the marmoset had to learn to open either of the two positions (left or right) to be rewarded. One session of 20 trials per day was conducted. The interval between sessions was aimed at 2-3 days (mean ± SD: UE, 2.50 ± 0.20; VPA, 2.54 ± 0.14: no statistical difference: p = 0.68). The inter-trial interval was 5 s. When 90% accuracy was reached in three consecutive sessions in Phase 1, the correct position was reversed on the next experimental session (Phase 2). When the marmoset reached the same criterion, the reward condition was reversed again (Phase 3). If the marmoset did not displace the cover for 60 s or opened the incorrect position cover, the trial was recorded as an error.

### Correlation across varieties of behavior tasks in UE and VPA-exposed marmosets

The results of each behavioral test were scored as follows in order to examine correlations. 1) social attention score were defined by the duration of attention toward the unfamiliar conspecific. 2) the inequity aversion score. Our previous study revealed that VPA-exposed marmosets failed to exhibit other-regarding behaviors (23). The unexposed marmosets refused to execute a simple task when they witnessed their partner receiving a more attractive reward for equal effort (inequity aversion). This inequity aversion was not observed in unexposed marmosets when the partner was absent. In contrast, VPA-exposed marmosets continuously executed the task, irrespective of their partners’ reward conditions. For this inequity aversion test, the ratio of food acceptance in the equity condition minus the ratio of food acceptance in the inequity condition was defined as the inequity aversion score. 2) the reciprocity evaluation score. The UE marmosets responded negatively to non-reciprocal human actors when they observed an unfair exchange between third parties who had no direct relevance to the marmosets, whereas the VPA-exposed marmosets did not exhibit any differential behavior between non-reciprocal and reciprocal conditions. For this third-party reciprocity test, the ratio of food acceptance from a reciprocator in the reciprocal condition to the ratio of food acceptance from a non-reciprocator in the non-reciprocal condition was defined as the reciprocity evaluation score.

### Statistical analysis

All statistical analyses were performed using JMP 16 software (SAS Institute, Cary, NC, USA). All values are expressed as the mean ± standard deviation, and p-values less than 0.05 were considered statistically significant.

## Results

### Three-chamber test

Marmosets underwent the three-chamber test five times during 15-19 weeks of age (childhood). Both the VPA-exposed and age-matched UE marmosets spent more time in the social chamber with an unfamiliar marmoset than in the other two chambers. However, there was no statistical difference among time spent in all three chambers (p=0.185, two-way repeated-measures ANOVA) in both control and model marmosets (Figure 2A). Marmosets in puberty underwent the three-chamber test once a week for five consecutive weeks between 29 and 41 weeks of age. As with the childhood results, there was no significant difference among the time spent in each chamber (p=0.921, two-way repeated-measures ANOVA, Figure 2B).

**Figure 2.**
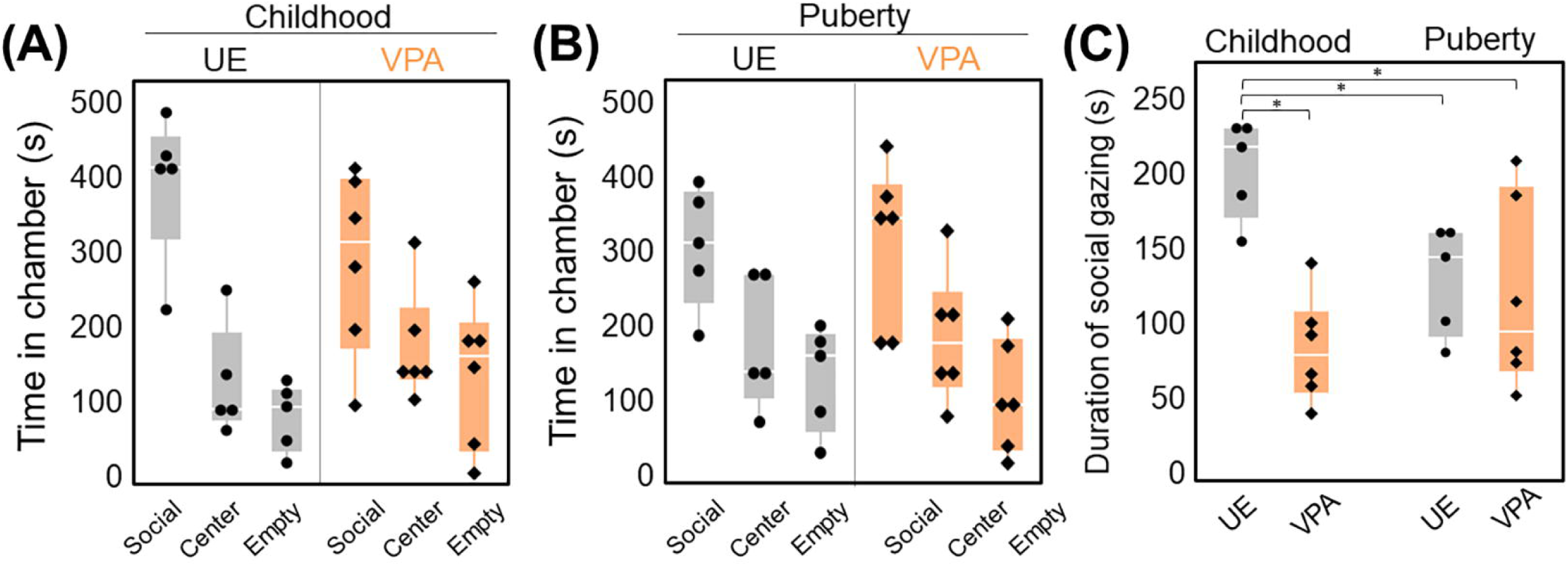
(A-C) Three-chamber test conducted both in childhood (15-19 weeks of age) and puberty (29-41 weeks of age). (A) Time spent in each chamber. There was a trend for a longer time spent in a social chamber in the unexposed (UE) group. Two-way repeated measure ANOVA, n=5 for UE, n=6 for valproic acid (VPA)-exposed. (B) Time spent in each chamber. There was no statical difference between the three chambers. Two-way repeated-measures ANOVA, n=5 for UE, n=6 for VPA. (C) The duration of social gazing in childhood in the UE group was significantly longer than that of the other three groups. Two-way repeated-measures ANOVA (F(1,9)=21.5525, p=0.0012) followed by Tukey’s HSD post-hoc test, n=5 for UE, n=6 for VPA. *: p<0.05.

We have further examined the duration of social gazing in marmosets in childhood and puberty. A two-factor repeated measures ANOVA showed a significant difference in the group factor (F(1,9)=7.3216, p=0.0242) and the interaction effects (F(1,9)=21.5525, p=0.0012). We did not find any significant difference in the age factor (F(1,9)=3.1127, p=0.1115). Multiple comparisons by Tukey’s post-hoc test revealed that the duration of social gazing in UE marmosets in childhood was significantly longer than that in VPA-exposed marmosets in childhood and in UE marmosets in childhood and puberty (Figure 2C).

### Position reversal learning

We have performed a serial reversal learning test to assess long-term preservation in the VPA-exposed marmoset (Fig. 3A). First, we examined the percentage of correct responses of the first 2-3 sessions in the first discrimination phase (Phase 1), the reversal phase (Phase 2), and the re-reversal phase (Phase 3; Figure 3A). The percentage of correct response of the first 2-3 sessions of the Phase1 was statistically significantly lager in the VPA-exposed marmoset than the UE marmoset (group factor (F(1,10)=8.0328, p=0.0177) and the interaction effects (F(1,10)=8.4483, p=0.0157: two-factor repeated measures ANOVA), and that of the Phase 3 was statistically significantly smaller (group factor (F(1,10)=6.0401, p=0.0338) and the interaction effects (F(1,10)=0.6950, p=0.4239: two-factor repeated measures ANOVA). The percentage of correct response of the first 2-3 sessions of the Phase 2 tended to be smaller in the VPA-exposed marmoset, although there was no statistical significance (group factor (F(1,10)=1.0478, p=0.3301) and the interaction effects (F(1,10)=0.8420, p=0.3804: two-factor repeated measures ANOVA). These results indicate that the VPA-exposed marmoset learn new skills more quickly, while they tend to adhere to previous learning.

**Figure 3.**
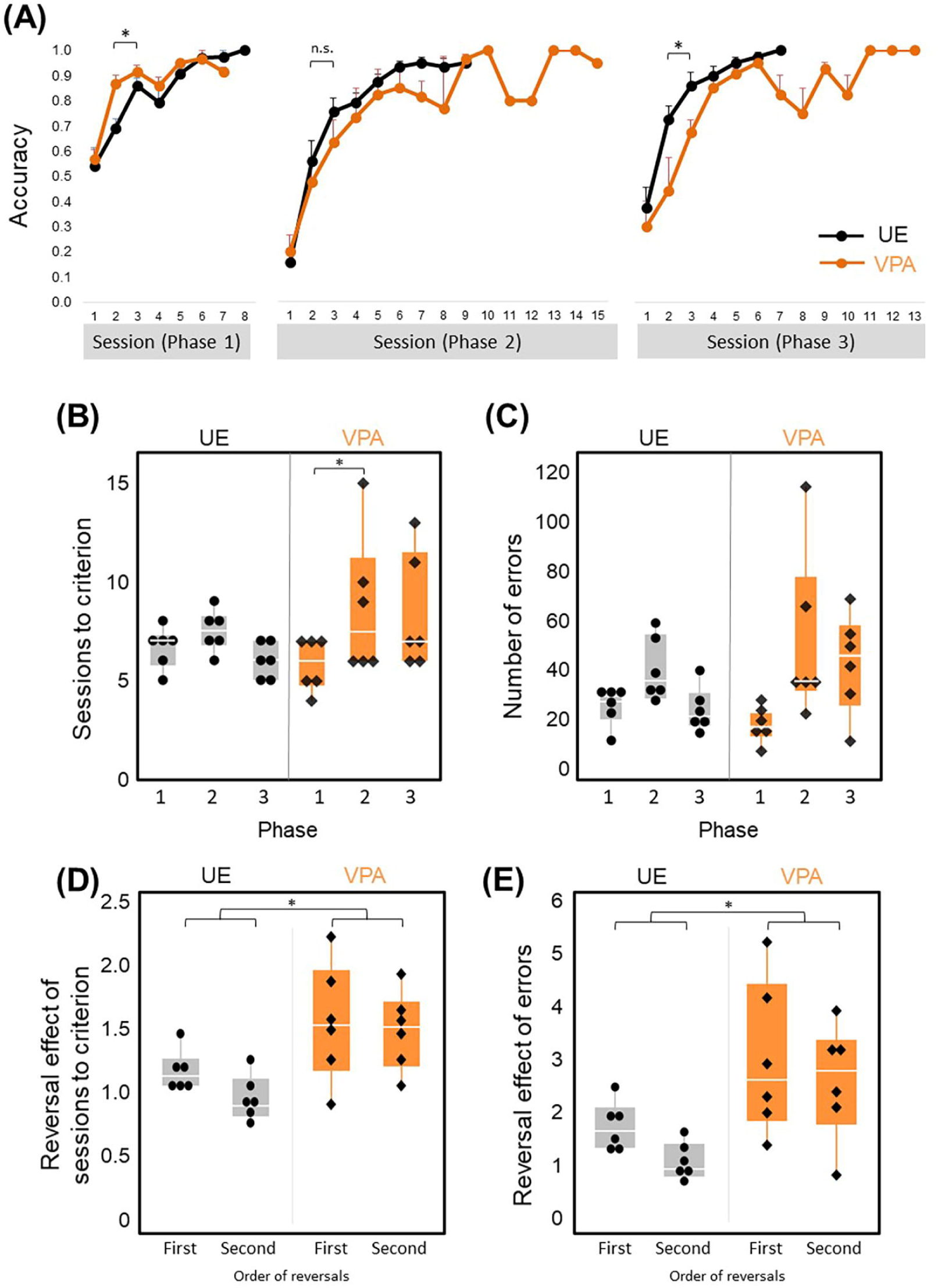
**(A-E)** Reversal Learning results across three phases. Phase 1 is acquisition trials. Phase 2 is reversal learning. Phase 3 is re-reversal learning. (n=6 for UE, n=6 for VPA) (A) The curve of the percentage of correct answers (±SE) across all trial phases. The performance of VPA-exposed marmosets was better for the first 2-3 days of phase 1 that UE marmoset, but worse for those of phase 2 and phase 3. However, statistically significant difference was observed only at Phase 1 and Phase 3. (B) Number of sessions required to achieve the criterion. The numbers of sessions of the second phase of the VPA marmosets were significantly larger than that of the first phase, while there was no difference between the numbers of the session of the UE marmosets. **(C)** Number of errors required for attainment criteria. (D) First and second reversal effects of number of sessions required to achieve the criterion. There is statistically significant group effect. (E) First and second reversal effects of the number of errors to reach the criterion. There is statistically significant group effect. *: p<0.05.

Second, we have examined the number of sessions and the number of errors to reach the criteria. A two-factor repeated measures ANOVA was performed to analyze the effect of VPA exposure group and learning phase on number of sessions to reach the criteria. The result showed a significant difference in the phase factor (F(2,20)=4.8271, p=0.0195) and the interaction effects (F(2,20)=3.6835, p=0.0435). We did not find any significant differences in the group factor (F(1,10)=0.8030, p=0.3913). Multiple comparisons by Tukey’s post-hoc test revealed that the number of sessions to reach criteria in phase 1 in the VPA-exposed marmoset was larger than that in phase 2 in the VPA-exposed marmosets (Fig. 3B). The numbers of errors to reach criteria produced the similar patterns (Fig. 3B).

A two-factor repeated measures ANOVA showed a significant difference in the phase factor (F(2,20)=10.5334, p=0.0008). We did not find any significant differences in the group factor (F(1,10)=0.9191, p=0.3603) and the interaction effects (F(2,20=3.1862, p=0.0629) (Fig. 3C). These results further supported response rigidity in model marmosets. Interestingly, both the average number of sessions and the average number of errors to reach the criteria (Fig. 3B, C) was smaller in Phase 1 in VPA-exposed marmosets than that in UE marmosets, although not statistically significant, adding evidence for higher performance in model marmosets in Phase 1.

Third, the number of sessions to reach the criteria in Phase 1 minus the number of sessions to reach the criteria in Phase 2 or 3, divided by the number of sessions reached in Phase 1, was defined as the first and second reversal effects, respectively (Fig. 3D, E). First and second reversal effects of the number of sessions to reach the criteria were larger in VPA-exposed marmoset than UE marmoset (group factor (F(1,10)=9.8345, p=0.0106) and the interaction effects (F(1,10)=0.6742, p=0.4307: A two-factor repeated measures ANOVA) (Fig. 3D). Furthermore, first and second reversal effects of errors to reach the criteria were also larger in VPA-exposed marmoset than UE marmoset (group factor (F(1,10)=8.4657, p=0.0156) and the interaction effects (F(1,10)=0.2204, p=0.6488: A two-factor repeated measures ANOVA). These results further endorsed the behavior perseverance of the VPA-exposed marmosets.

### Correlations across varieties of behavior tasks in UE and VPA-exposed marmosets

Social attention score in UE and VPA-exposed marmosets during childhood in the three-chamber task correlated with the behavioral scores of inequity aversion (23) in adults (r=0.879, p=0.0092; Figure 4A). These short attention scores were correlated with the discrimination scores of the third-party evaluation test (22) in adults (r=0.943, p=0.0004; Figure 4B). The second reversal effect in adults was correlated with social attention score during childhood (r=0.879, p=0.0092; Figure 4C). There was also a strong correlation between inequity avoidance and third-party reciprocity assessment scores (r=0.920, p=0.0092, Figure 4D). The strong correlations among the scores of three social tests (social attention during childhood, inequity avoidance, and third-party reciprocity in adulthood) endorse the robust social impairment of VPA-exposed marmosets. In addition, the second reversal effect score was correlated with inequity aversion (r=0.779, p=0.0134; Figure 4E). The reciprocity evaluation score tended to correlate with the second reversal effect although this was not statistically significant (r=0.791, p=0.0610; Figure 4F). The social attention score during puberty was not correlated with any of the other behavioral tests performed (Table 1).

**Table 1.** Pearson correlation coefficient r between the duration of social gazing in childhood and puberty, the reciprocity evaluation score, the inequity aversion score and the second reversal effect of reversal learning test. P-values indicate statistical significance (*p<0.05, **p<0.01, and ***p<0.001).

|  | Duration of<br>social gazing<br>-childhood- | Duration of<br>social gazing<br>-puberty- | Reciprocity<br>evaluation score | Inequity<br>aversion score | Second<br>reversal effect |
| --- | --- | --- | --- | --- | --- |
| Duration of<br>social gazing<br>-childhood- |  | 0.413 | 0.943 | 0.879 | 0.735 |
| Duration of<br>social gazing<br>-puberty- |  |  | 0.319 | -0.047 | 0.284 |
| Reciprocity<br>evaluation<br>score | *** |  |  | 0.920 | 0.791 |
| Inequity<br>aversion score | ** |  | ** |  | 0.779 |
| Second<br>reversal effect | * |  |  | * |  |

\*p<0.05, \*\*p<0.01, \*\*\*p<0.001

**Figure 4.**
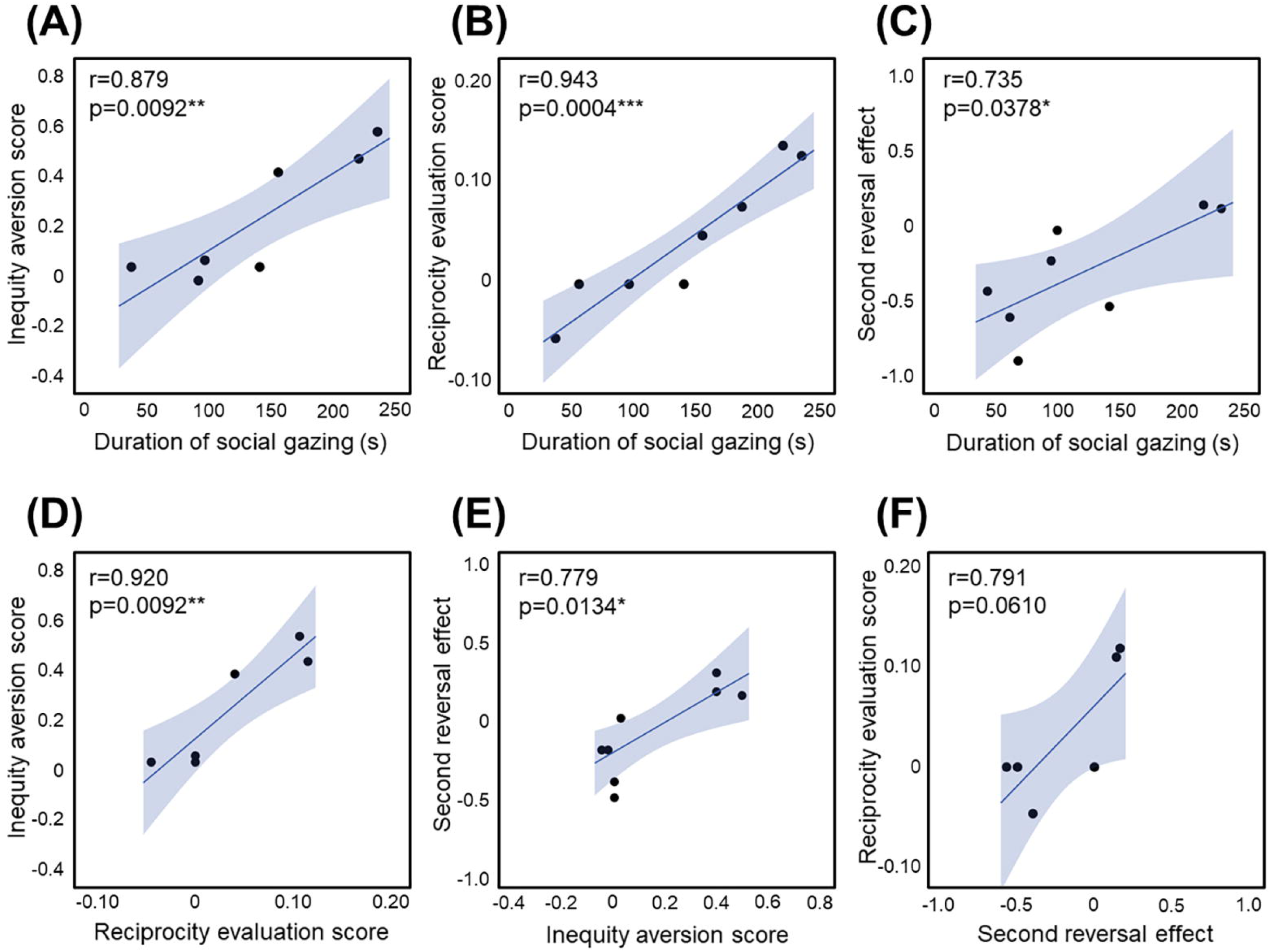
(**A**) Correlation between the inequity aversion score and the duration of social gazing in childhood (n=3 for UE, n=4 for VPA). (**B**) Correlation between the reciprocity evaluation score and the duration of social gazing in childhood (n=4 for UE, n=4 for VPA). (**C**) Correlation between the second reversal effect and the duration of social gazing in childhood (n=2 for UE, n=6 for VPA). (**D**) Correlation between the inequity aversion score and the reciprocity evaluation score (n=3 for UE, n=3 for VPA). (**E**) Correlation between the second reversal effect and the inequity aversion score (n=5 for UE, n=4 for VPA). (**F**) Correlation between the reciprocity evaluation score and the second reversal effect (n=2 for UE, n=4 for VPA). *p<0.05, **p<0.01, and ***p<0.001.

## Discussion

In the present study, we showed that the VPA-exposed marmoset spent less time gazing at other individuals than the UE marmoset in childhood. This short social attention time in childhood was well correlated with impairment of two higher-order social abilities (inequity aversion and third party reciprocity) found in adulthood in the same animal (22, 23). These results are consistent with the notion that early deficit of social gazing enhance later ASD symptom (11,12,37). Furthermore, the reversal learning test revealed adherence to previous long-term learning in VPA-exposed marmosets in adulthood, which was also associated with shorter gaze time to other individuals in childhood.

Gaze disruption to other conspecifics in VPA-exposed marmosets during childhood is consistent with studies of human children with ASD using home movies. Up to 6 months of age (9) and from 12 to 30 months of age (8), infants diagnosed with ASD spend less time looking at and orienting toward people than normal children. Also, Many previous studies using eye-tracking systems have found that children and individuals with ASD spend less time looking into the eyes of others (10,12,38). However, in the present study, there was no difference in social attention between UE marmosets and VPA-exposed marmosets in puberty. This may be because social gazing in UE marmosets become shorter in puberty than in childhood (Fig. 2C), making the differences between the groups too small to detect. Adult marmosets share food items with marmosets in childhood (10-18 weeks of age), but after puberty (24 weeks of age), the probability of sharing food is less than 1/7 of that in childhood (39). Reduced reward values in pubertal marmosets may be associated with reduced gaze duration. In the next step, it would be worthwhile to investigate the degree of attention paid to the eyes in the UE and VPA-exposed marmosets in a sophisticated manner throughout their lives. An eye-tracking system revealed social target avoidance in adulthood in two types of ASD model macaques induced by immune activation and exposure to VPA *in utero* (40,41) and showed social feature preference in *MeCP2* mutant ASD model macaques in subadults (42). However, the gaze of these ASD model monkeys has not yet been examined in childhood. There are no studies of social attention by gaze in rodent models.

In young children (12-24 months) with ASD, abnormalities in eye contact have been correlated with deficits in empathy and prosocial behavior in preschool (43). Longitudinal studies have shown that joint attention is associated with social cognitive outcomes such as language development (44) and Theory of Mind (45). Early delays in basic face processing, including attention to faces, have been considered to contribute to an atypical trajectory of social communication skills in individuals with ASD (11). Weak attention to others in early childhood in marmosets examined in this study was correlated with performance on two social cognitive tasks in adulthood. This result provides evidence that the behavioral phenotype of this animal model is robust as a social disorder and is consistent with the notion that weak attention to others in early childhood amplifies the trajectory of the development of autistic symptoms later in life (11,12,37,46).

The VPA-exposed marmosets showed a quicker increase in performance during the first discrimination phase (Phase 1), and tended to require fewer sessions and fewer errors before reaching the criteria. The personal strength of memory is often seen in children with ASD (48). For example, superior performance for individuals with high functioning ASD was frequently reported in tasks involving recall of a path in maze, and shorter learning times in a map learning task (52). They also can remember places they have been once and remember exactly where things are located. The higher capability of the VPA-exposed marmoset in Phase 1 suggested that the failure of the VPA-exposed marmoset in social tasks previously reported (22,23) cannot be ascribed to a decline in intellectual ability.

The VPA marmoset showed perseveration, as measured by the number of sessions to reach the learning criterion in reversal phases and by the first and second reversal effects. A slower increase in the percentage of correct responses in Phase 2 and 3 also supported response rigidity in the VPA-exposed marmoset. Consistent with this phenotype of the VPA-exposed marmoset, individuals with ASD persist in habits and sameness and resist changes in the environment and their behaviors (47). Repetitive habits become part of their routines in people with ASD. A previous showed that the VPA marmosets in childhood show a call pattern with low entropy that is biased toward one type of call (phee call) (24). Similarly, the involuntary repeat of words and phrases (palilalia) is frequently observed in people with ASD (53). These results suggest that the VPA-exposed marmoset both in childhood and adulthood is primed for restricted and repeated behavior, one of the two core symptoms of ASD.

Gazing time scores in childhood and social skill scores in adulthood in the marmosets examined here correlated with adult rigidity scores, suggesting that the behavioral phenotypes exhibited by VPA-exposed marmosets may be rooted in the same pathophysiology of the brain. The medial prefrontal cortex has been implicated in sociality and cognitive flexibility (49–51). Serotonin depletion in the prefrontal cortex severely impairs the reversal learning performance (34). Interestingly, downregulation of the serotonin receptor-encoding gene *HTR5A* has been reported in the cortex of VPA-induced marmosets (24).

## Conclusion

Results in this study suggest that the VPA-exposed marmoset will provide a useful platform for investigating the impact of early rectification of weak social attention on the progression of ASD symptom, and will help for developing effective early treatments. Previous synaptic and gene expression studies of the VPA-exposed marmoset have indicated that the brains of human children with ASD may exhibit aberrant plasticity and critical period (24). Atypical plasticity can alter the trajectory of the normal development of social brain circuits, even if social experiences are normalized by behavioral interventions. The administration of drugs that correct anomalous circuit plasticity combined with behavioral interventions may further enhance the effectiveness of early treatment.

## Limitation

In this study, we could not evaluate the gaze direction of the conspecifics in the cylinders because their heads were often not visible, even from the three video cameras. Eye contact, or joint attention, is the exchange of gaze between two individuals, and its impairment is a major symptom of ASD. As marmosets are a rare species among non-human primates that engage in eye contact (16), it would be valuable to monitor the gaze of both animals and partnered unfamiliar conspecifics in the future.

To reduce the stress of the young marmoset, we did not conduct a controlled experiment to confirm that impaired head orientation to the conspecifics of the VPA-exposed marmoset reflected an attention disorder rather than a physical defect. For example, instead of an adult marmoset, an experiment in which an object was placed in the cylinder was considered. However, the strong correlation between scores for head orientation to other individuals in childhood and scores for social experiments in adulthood supports the existence of impaired social attention in young VPA-exposed marmosets.

## Acknowledgments

We thank Ms. Akiko Tsuchiya for providing technical support.

This work was supported by an Intramural Research Grant (Nos. 23-7, 26-9, and 29-6 to N.I.) for Neurological and Psychiatric Disorders from the NCNP, KAKENHI (Nos. 16H02058 and 15K13159 to N.K, 25882032 to M.Y., 2460020 and 15K01791 to A. N.), and AMED Grant Number JP21dm0207066 (NI).

## Conflict of Interest

The authors declare that the research was conducted in the absence of any commercial or financial relationships that could be construed as potential conflicts of interest.

## Author Contributions

N.K. and N.I. designed this study. A.N. and M.Y. performed behavioral experiments. N. A., M. Y., and M. N. analyzed the data. K.N. managed the production and physical condition of the animals. N. K., N. I., A. N., M. Y., and M. N. wrote the manuscript. All authors have read and approved the final manuscript.

## Notes

### Competing Interest Statement

The authors have declared no competing interest.

